# Minimal-invasive enhancement of auditory perception by terahertz wave modulation

**DOI:** 10.1101/2021.06.01.446562

**Authors:** Xiaoxuan Tan, Kaijie Wu, Shuang Liu, Chao Chang, Wei Xiong

## Abstract

Hearing impairment is a common disease affecting a substantial proportion of the global population. Currently, the most effective clinical treatment for patients with sensorineural deafness is to implant an artificial electronic cochlea. However, the improvements to hearing perception are variable and limited among healthy subjects. Moreover, cochlear implants have disadvantages, such as crosstalk derived from the currents that spread into non-target tissue between the electrodes. Here, in this work, we describe terahertz wave modulation (THM), a new, minimally invasive technology that can enhance hearing perception in animals by reversible modulation of currents in cochlear hair cells. Using single-cell electrophysiology, guinea pig audiometry, and molecular dynamics simulations (MD), we show that THM can reversibly increase mechano-electrical transducer (MET) currents (∼50% higher) and voltage-gated K^+^ currents in cochlear hair cells through collective resonance of −C=O groups. In addition, measurement of auditory brainstem response (ABR) in guinea pigs treated with THM indicated a ∼10 times increase in hearing sensitivity. This study thus reports a new method of highly spatially selective hearing enhancement without introducing any exogeneous gene, which has potential applications for treatment of hearing disorders as well as several other areas of neuroscience.

## Introduction

Auditory perception is a complex process involving a highly specific information transmission process, with cochlea playing a vital role in the processing of sound information in animals (Sun et al., 2018; Qiu et al., 2018). The cochlea transmits sound through mechanical changes to convert mechanical energy into electrical energy, ultimately generating nerve impulses, which are then transmitted to the central system to produce hearing (Clause et al., 2014). Hearing impairment or deafness, such as sensorineural hearing loss, is generally caused by damage to the hair cells of the inner ear (Corns et al., 2018; Wu et al., 2016; Delmas et al., 2013).

At present, treatment for hearing impairment is primarily administered through pharmacological treatment, hearing aid equipment, stem cell differentiation and transplantation (Oshima et al., 2010; Li et al., 2003; Chen et al., 2012), optogenetics (Huet et al., 2021), and electronic cochlear implantation (Wilson et al., 1991; Kipping et al., 2020; Gang et al., 2008). For patients with severe deafness, the most effective treatment is to surgically implant an electronic cochlea. However, the utility of artificial electronic cochleas is severely reduced due to the current diffusion effect, since the number of electrodes that can be implanted is very limited, while too few electrodes will lead to low-frequency resolution (Lee et al., 2001; Friesen et al., 2016). Development of stem cell differentiation and transplantation therapeutic strategies has been hindered by ethical considerations, and optogenetics requires expression of exogenous genes, and is thus a currently unsuitable approach for human patients.

Near-infrared triggering of cochlear hearing has been also developed as a viable potential method to restore hearing. This technique relies on the use of relatively short wavelength, near-infrared (NIR) light stimulation in short pulses to induce optically-evoked auditory brain response (OABR) in the cochlea (Wang et al., 2016; Wang et al., 2015). However, the underlying mechanism for the OABR is still poorly understood (Richard et al., 1971), and does not evoke the patterns of neural excitation that are normally associated with acoustic stimulation. Here, we introduce a different hearing enhancement technology, Terahertz wave modulation (THM), a minimally invasive technique for transmitting terahertz wave energy to the cochlea which subsequently improves the sensitivity of the cochlear outer hair cells and the K^+^ conduction velocity. This technology can increase the hearing threshold of animals with good spatial selectivity, no stimulus artifacts, and no current diffusion effect (Izzo et al., 2007; Izzo et al., 2006). We further identify several significant advantages of optical nerve stimulation nerves over electrical stimulation, particularly in the application of more spatially confined neural stimulation.

In this study, we performed whole-cell patch clamp recording of the outer hair cells on the basilar membrane of acutely isolated mouse cochleas. We found that optical terahertz wave modulation (THM) at a wavelength of 8.6 μm (34.88 THz) can be used as a quasi-non-invasive means of enhancing hair cell sensitivity. We show that THM can significantly increase both K^+^ and MET currents of the cochlear outer hair cells, and that MET currents exhibit a positive correlation between enhancement amplitude and irradiation power. As a striking example, we found THM can increase the hearing threshold of adult guinea pigs by approximately 10 dB. We also confirmed by kinetic simulation that this effect may be attributed to the characteristics and kinetic changes in K^+^ channels. These changes are likely due to THM-induced promotion of channel transport and regulation of the nature of the channel subunits that form pores, causing a collective resonant vibration of the −C=O bond in the filter component of the ion channel.

## Results

### THM significantly enhances MET currents

THz fibers (core diameter 600 μm, NA 0.35) deliver THz light energy to the target area of the isolated cochlea (Fig. 1A). Cochlear hair cells convert sound stimuli into electrical signals through gating by mechanically sensitive ion channels in their stereociliary (hair) bundle. The MET channels are thus essential for maintaining normal function of the cochlea (Corns et al., 2018). Therefore, we used a fluid jet stimulation system to detect the MET currents following a previously used method in which the directions of positive and negative polarity are reversed (Zhao et al., 2014; Richter et al., 2008). MET currents were recorded in one-minute cycles, with THM applied at 10s and then switched off at 40s (Fig. 1C). During THM treatment, the MET currents increased significantly (628.8±94.22 vs 686.7±89.08 pA, n=8, t_7_=3.699, *P*=0.0077, paired Student’s *t*-test), then stabilized after approximately 5s (Fig. 1, D and E). When THM was removed at 40s, the MET currents immediately decreased (686.7±89.08 vs 608.5±79.56 pA, n=8, t_7_=3.366, *P*=0.0120, paired Student’s *t*-test), and again stabilized after about 5s (Fig. 1, D and E). We detected no significant difference between the MET channel currents during the first ten seconds prior to treatment and the last ten seconds after removing treatment (Fig. 1E, 628.8±94.22 vs 608.5±79.56 pA, n=8, t_7_=0.5771, *P*=0.5820, paired Student’s *t*-test), which indicates that the effect is reversible.

**Figure 1.**
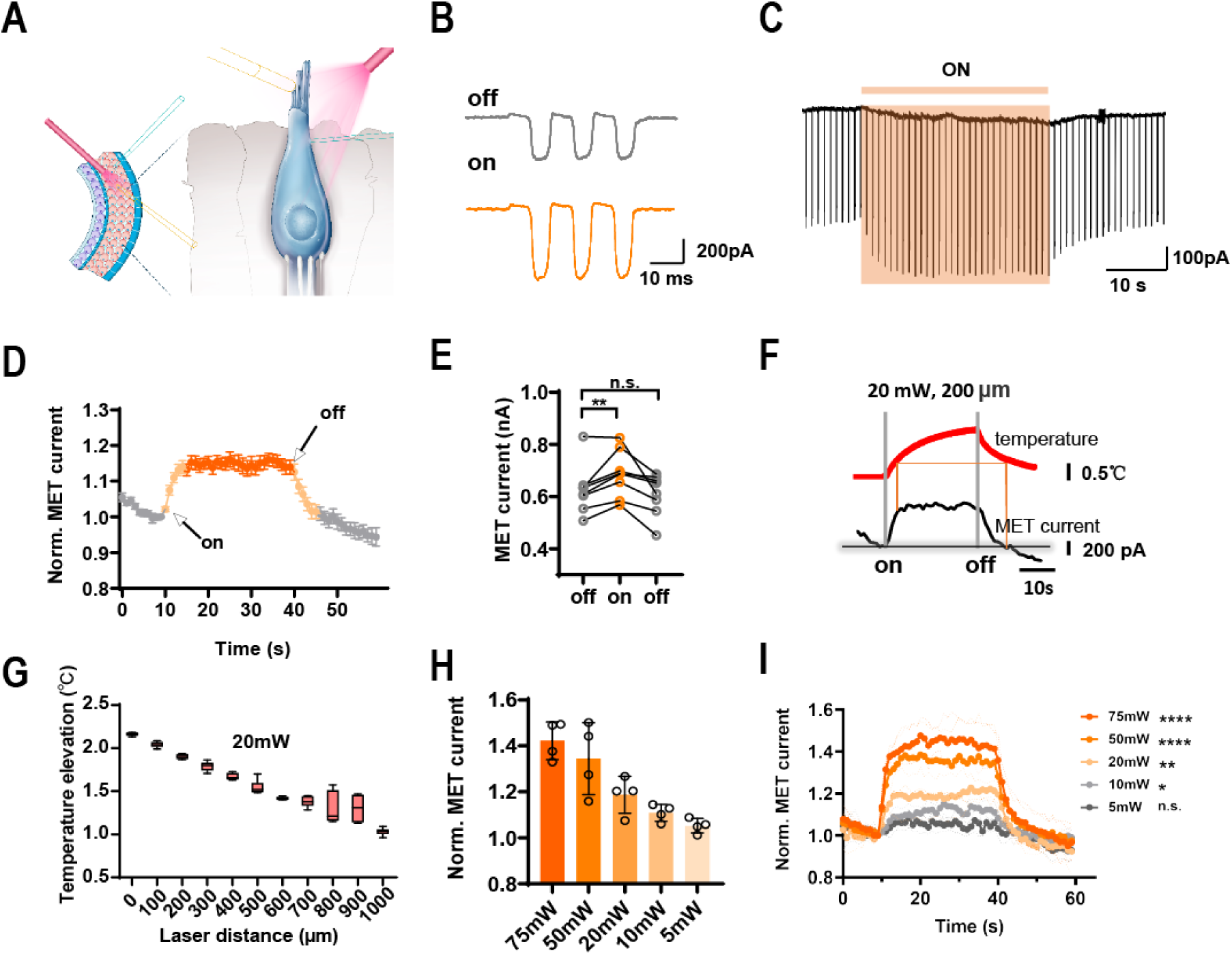
Terahertz wave modulation (THM) enhances MET currents. **(A)** Schematic of whole-cell recording configuration from OHCs. **(B)** MET currents in control (gray) and during THM (orange). **(C)** THM increases MET currents (shaded part). **(D)** MET currents change with time, orange indicates the THM period (including light orange and dark orange). **(E)** Comparison of MET currents before, during and after THM (n=8 cells). **(F)** Schematic diagram of temperature and MET currents over time. **(G)** showing temperature elevation with respect to the distance from the fiber tip (n=6). **(H-I)** The influence of different powers of THM on MET currents (currents are normalized, n=4 cells). Paired Student’s *t*-test, n.s. *P* >0.05, * *P* <0.05,** *P* <0.01, \*\*\*\**P*<0.0001.Error bars represents s.e.m.

Even though we chose a wavelength (8.6 μm) with a low attenuation band (Liu et al., 2021), water still has a high terahertz absorption rate, which may produce thermal heat. Previous studies have shown that heating can enhance or inhibit the activity of neurons (Albert et al., 2012; Stujenske et al., 2015), so we tested the temperature rise in the extracellular fluid at different locations from the optical fiber (Fig. 1G). The cell under test is about 200 μm from the fiber tip, after 5 seconds of irradiation, the temperature at this location increased by about 0.5°C and continued to rise, but at this time, the MET currents had risen to near the maximum value and remained stable. In addition, when the temperature increases by the same value (the two points where the orange line intersect with the temperature curve), the MET currents varies significantly, with one near the maximum value and the other returning to the pre-irradiation level (Fig. 1F). The relationship between temperature and current was strikingly nonlinear, suggesting temperature-independent modulation of a MET current.

Next, we tested the power dependency of THM (responses to different power levels were measured within individual cells, n=4), and we also tested the temperature variation at different power levels (fig. S2). we found that the peak of MET currents increased as the power of the light source increased (Fig. 1H). At 75 mW (the maximum power of our laser), the MET currents increased by approximately 50% (Fig. 1I). We next sought to determine whether increases in current were due to cumulative effect or increases in power. We found that the current also decreased with decreased power of THM stimulation, therefore allowing us to exclude cumulative effects. These findings indicated that THM enhances the activity of MET channels and the sensitivity of hair cells, thus exerting a strong regulatory effect on the response of hair cells to external mechanical stimuli, with a positive correlation between the amplitude of current enhancement and THM intensity.

### THM modulates voltage-gated K^+^ channels

K^+^ currents are the most important cochlear conductive currents caused by cochlear potential (Jovanovic et al., 2020; Dierich et al., 2020; Babola et al., 2020). Hair cell K^+^ channels play a key role in regulating hair cell excitability and cochlear potential transduction (Wang et al., 2015; Johnson et al., 2011; Kang et al., 2017). Therefore, we tested whether THM has a regulatory effect on hair cell K^+^ channels (Fig. 2A). To this end, we performed patch clamp recordings of hair cells in which we increased the command voltage from -150 mV to 110 mV in increments of 20 mV to measure the activation of K^+^ and Na^+^ channels (Fig. 2, B and C). The results showed that Na^+^ currents increased, but only slightly and non-significantly, indicating that the effect of THM on Na^+^ currents is negligible.

**Figure 2.**
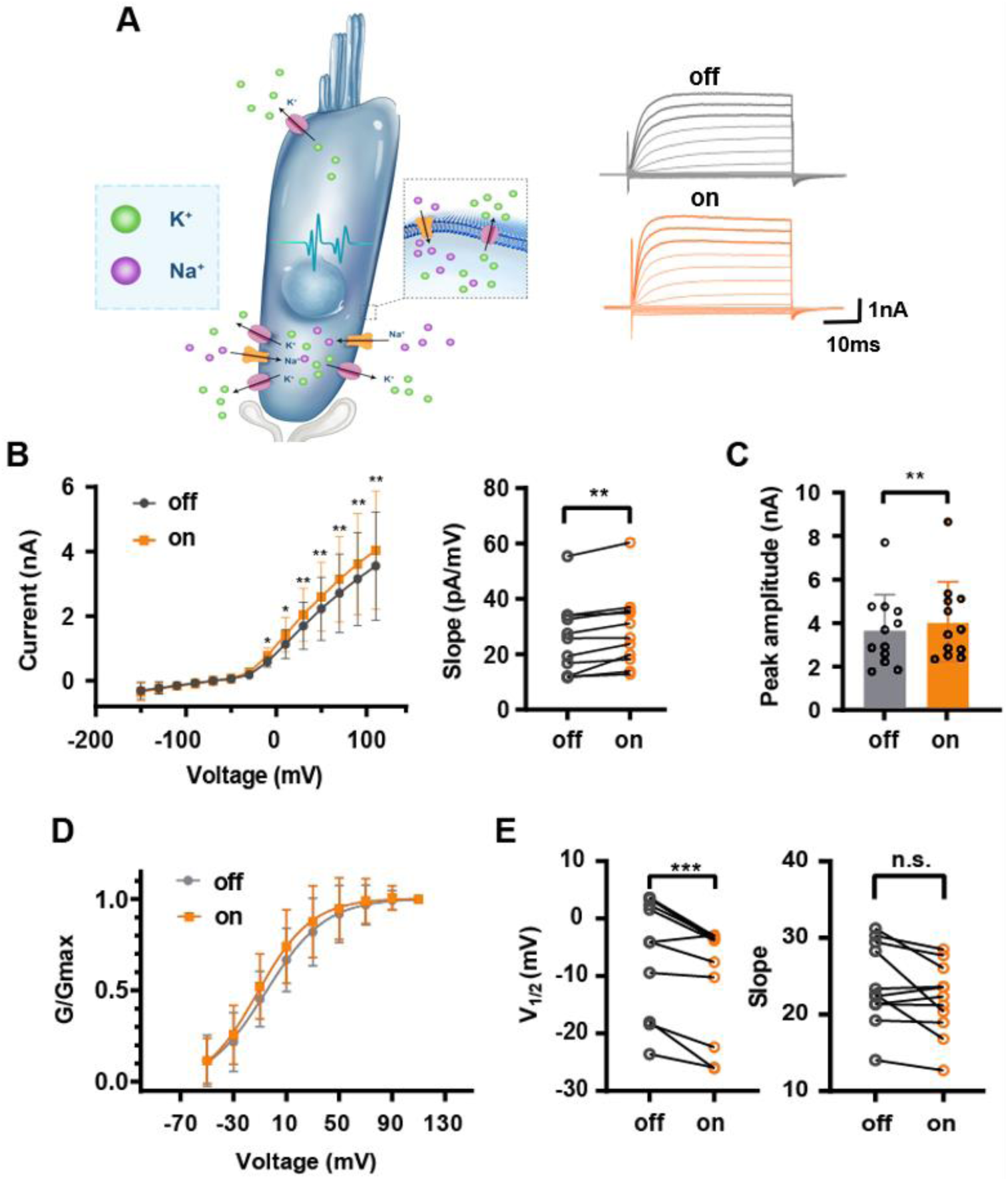
THM enhances K ^+^ currents. (**A**) Left, distribution of ion channels in hair cells. Right, MET currents in control (gray) and during THM (orange). (**B**) Left, I-V curves. Right, comparison of the I-V curve slopes (n=11 cells). (**C**) Peak amplitudes of K ^+^ currents (n=12 cells). (**D**) Activation curves of the K^+^ currents (n=11 cells). (**E**) Group data comparing the half-activation voltages and the slopes (n=11 cells). n.s. *P* >0.05, \**P* <0.05, \*\**P* <0.01, \*\*\**P* <0.001, **** *P* <0.0001, Paired Student’s *t*-test and repeated measures ANOVA with post-hoc Bonferroni’s multiple comparison. Error bars represents s.e.m.

In contrast, we observed a significant increase in the amplitude of K^+^ currents during THM (3552±1676 vs 4035±1829 pA, n=12, t_11_=3.976, *P*=0.0022, paired Student’s t-test, Fig. 2C), with the slope of the IV curve increasing from 25.52±13.29 to 28.46±13.62 pA/mV (n=11, t_10_=4.335, *P*=0.0015, paired Student’s t-test, Fig. 2B). Interestingly, THM shifted the activation curve toward hyperpolarized V_m_ by about 5 mV (V_1/2_: -5.714±10.13 vs -10.21±9.683 mV, n=11, t_10_=5.090, *P*=0.0005, paired Student’s t-test, Fig. 2, D and E), although there was no significant change in the slope of the activation curve (23.94±5.298 vs 21.97±4.715, n=11, t_10_=2.164, *P*=0.0557, paired Student’s t-test, Fig. 2E). These findings show that THM can enhance the activity of K^+^ channels, thereby increasing the rate of K^+^ transport. In summary, our results indicate that THM has a strong regulatory effect on hair cell responsiveness and cochlear conduction current, which is likely attributable to THM-associated changes in the characteristics of voltage-gated K^+^channels.

### THM causes collective resonance in the −C=O groups of K^+^ channels

To explore the mechanisms underlying the molecular dynamics driving ion current enhancement by THM, we constructed a model ion channel with a highly selective filter (fig. S3). Using this model, we found that THM accelerates the speed of K^+^ permeation (*i.e.,* super-permeation) through the K^+^ channel, but has no significant effect on Na^+^ channels (Fig. 3A). In general, ion permeability is determined by the number of ions that can penetrate from one side of the channel to the other (Wojciech et al., 2018). As shown in Fig. 3B and C, a comparison of permeation events between K^+^ to Na^+^ (∼4:1) through the K^+^ channel and Na^+^ to K^+^ (∼3:1) through the Na^+^ channel demonstrates the high selectivity of ion permeation in the absence of THM.

**Figure 3.**
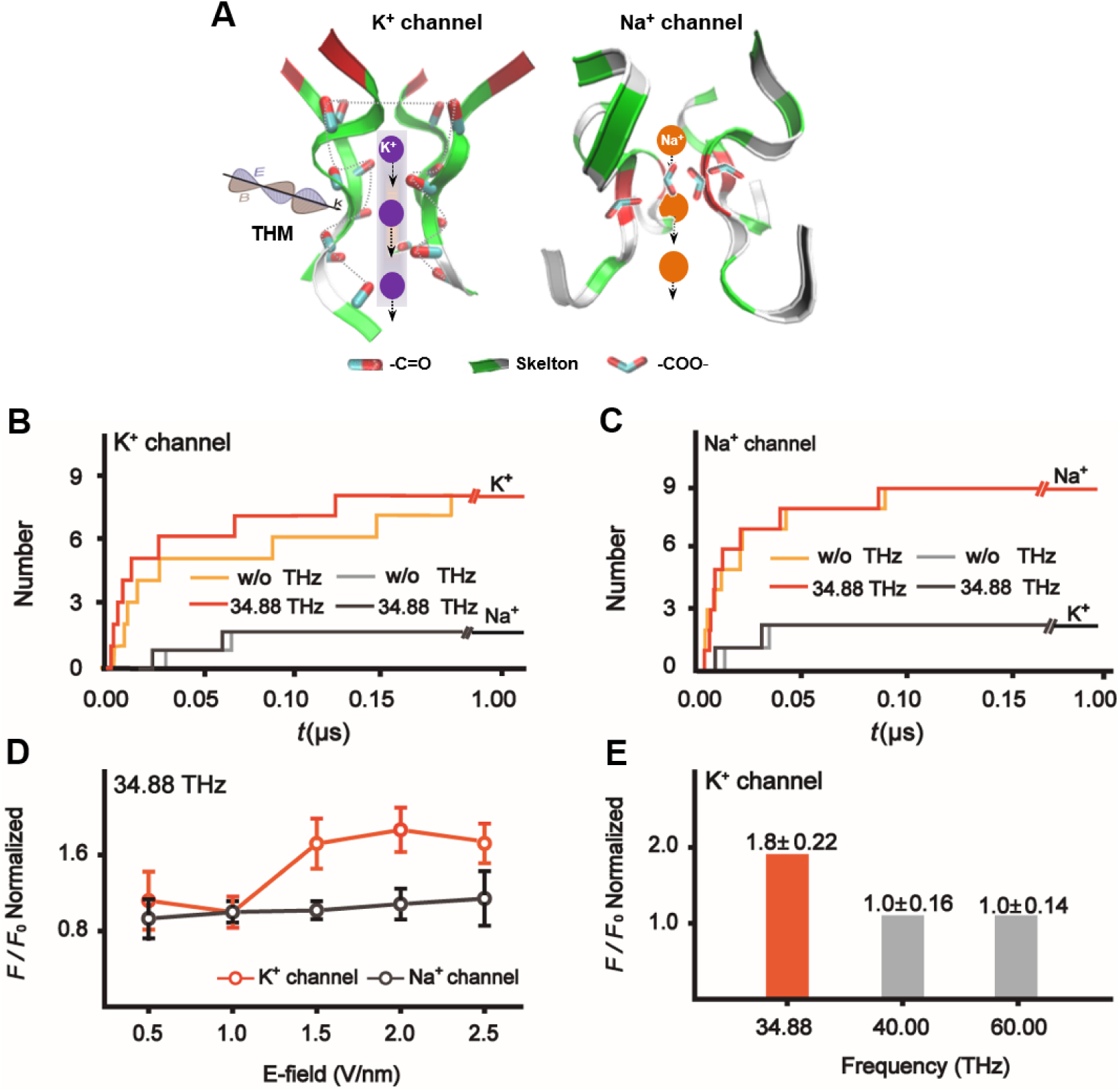
The discrepancy between K^+^ and Na^+^ permeation without and with THM. (**A**) Schematic representation of the structure of filter region for K^+^ channel and Na^+^ channel interaction with THz wave. The dotted line denotes THz waves are absorbed by K^+^ channels, but not by Na^+^ channels. The number of K^+^ (orange and lighter orange curves) and Na^+^ (black and lighter black curves) permeate through K^+^ channel (**B**), through Na^+^ channel (**C**), with and without THM at frequency *γ=*34.88 THz, and strengthen *A*=2.5 V/nm, respectively. (**D**) The ratio of permeation of K^+^ and Na^+^ (*F* / *F*_0_) depending on the THM intensity for K^+^ channel (orange curves) and Na^+^ channel (black curves). (**E**) The ratio of the permeation of K^+^ and Na^+^ (*F* / *F*_0_) dependent on the frequency of THM for K^+^ channel.

Notably, the average speed of K^+^ translocation is ∼2-fold greater through the K^+^ channel under THM treatment (Fig. 3B), but does not obviously change during THM treatment of the Na^+^ channel (Fig. 3C). Furthermore, we defined the average ratio of ion translocation speed (*F*/*F*_0_) to explore the difference in ion permeation attributable on THM. In Fig. 3D, we further found that the *F*/*F*_0_ of K^+^ was markedly accelerated for the K^+^ channel, but remained unchanged for the Na^+^ channel with increasing THM strength. However, we also observed that the *F*/*F*_0_ of K^+^ permeation was increased by THM solely at *γ*=34.88 THz, but retained lower efficiency at frequencies of *γ*=40.0 and 60.0 THz, thus indicating that K^+^ super-permeation is frequency-dependent (Fig. 3E). Together, these results reveal that THM at *γ*=34.88 THz resulted in K^+^ super-permeation through the K^+^ channel, but not the Na^+^ channel.

To explain the molecular mechanisms responsible for differences in the effects of THM on K^+^ and Na^+^ channels, we considered the dependence of physical properties of carbonyl (−C=O) groups of the K^+^ channel and carboxyl (−COO^-^) groups of the Na^+^ channel on THM frequency. Fig. 4A shows a remarkably strong absorption finger located in the range of absorption half-height width (FWHM) (∼2.0 THz) near that of 34.88 THz for the K^+^ channel, while the absorption finger of −COO^-^ groups are distant from 34.88 THz for the Na^+^ channel. These vibration modes of absorbed dipoles are illustrated in Fig. 4A. These findings suggest that THz waves are perfectly absorbed by K^+^channels, but not by Na^+^ channels. Furthermore, we identified the absorption modes of −C=O groups for the K^+^ channel using the density functional (DFT) method. We found that the finger of these connections, excluding the −C=O groups, deviated from 34.88 THz, thus indicating that 34.88 THz mainly contributes to the vibration of −C=O groups, but not to the vibration of –N–H– groups (in the backbone of filter region) (Fig. 4B). It warrants mention here that 34.88 THz subjects a total of 16 pairs of −C=O groups to co-oscillation (fig. S4). Therefore, we next calculated the vibrations of K^+^ together with the −C=O groups by solving the phonon vibration equation (1) as follows (Huang et al., 2014),

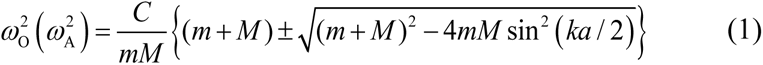

**Figure 4.**
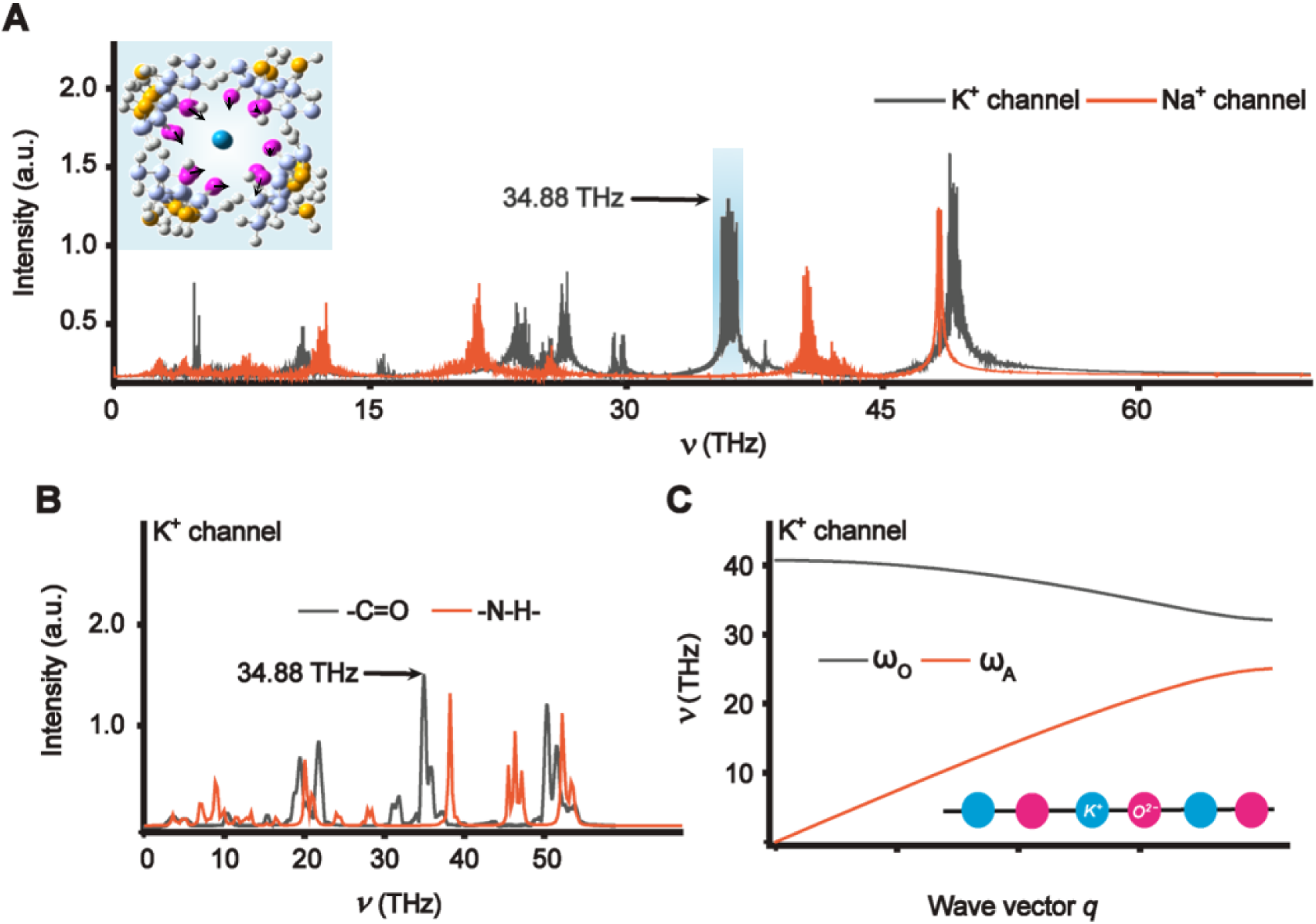
The super-permeation mechanism on the selective absorption of THM for ion channel. (**A**) The absorption spectra of –C=O groups for K^+^ channel and –COO-groups for Na^+^ channel, respectively, calculated by using the MD method, and the overall collective vibration modes of –C=O groups in the filter of K^+^ channel is shown in the illustration. The region of the blue stripe represents that 34.88 THz falls in half-width of the absorption finger. (**B**) Calculations on the eigen mode of –C=O groups and the –N–H– connections (excluding −C=O groups of the skeleton of the filter region) for K^+^ channel by using density functional (DFT) method. (**C**) The phonon frequency of K^+^ at internal binding sites participating in entirety collective vibrations of –C=O groups for K^+^ channel calculated by density functional (DFT) method. The interaction of K^+^ and the filter of K^+^ channel is simplified to form the one-dimensional (1D) phonon chain (O^2-^-K^+^-O^2-^-K^+^). The phonon mode ω_o_ and ω_A_, respectively, denote the vibration of O^2-^ and K^+^ in distinct and the same directions.

where *M* and *m* indicate the masses of O^2-^ and K^+^, respectively; phonon modes*ω*_O_ and*ω*_A_ respectively denote the vibrations of these ions in distinct or the same directions; *C* represents the coupling constant. Our results demonstrate that the vibrations of K^+^ vary in the range of 32 ∼ 40 THz (Fig. 4C), showing that 34.88 THz corresponds to the participation of K^+^ in the collective oscillations of the −C=O groups. Therefore, the THz wave absorbed by the −C=O groups of K^+^ channels leads to their collective resonance, thereby enhancing super-permeation by K^+^ ions, whereas this absorption and subsequent super-permeation cannot occur in Na^+^ channel.

### THM enhances the hearing perception of guinea pigs

We next tested whether THM has any effect on the hearing threshold *in vivo*. Considering that the cochlear structure of guinea pigs is highly similar to that of humans, we conducted these experiments using guinea pigs. As shown in Fig. 5A, the guinea pig’s posterior ear window is opened to expose the cochlea, and THM is delivered approximately 1 mm away from the cochlear oval window. The light spot size was just adequate to cover the guinea pig oval window (a circle with a diameter of approximately 2 mm, fig. S1).

**Figure 5.**
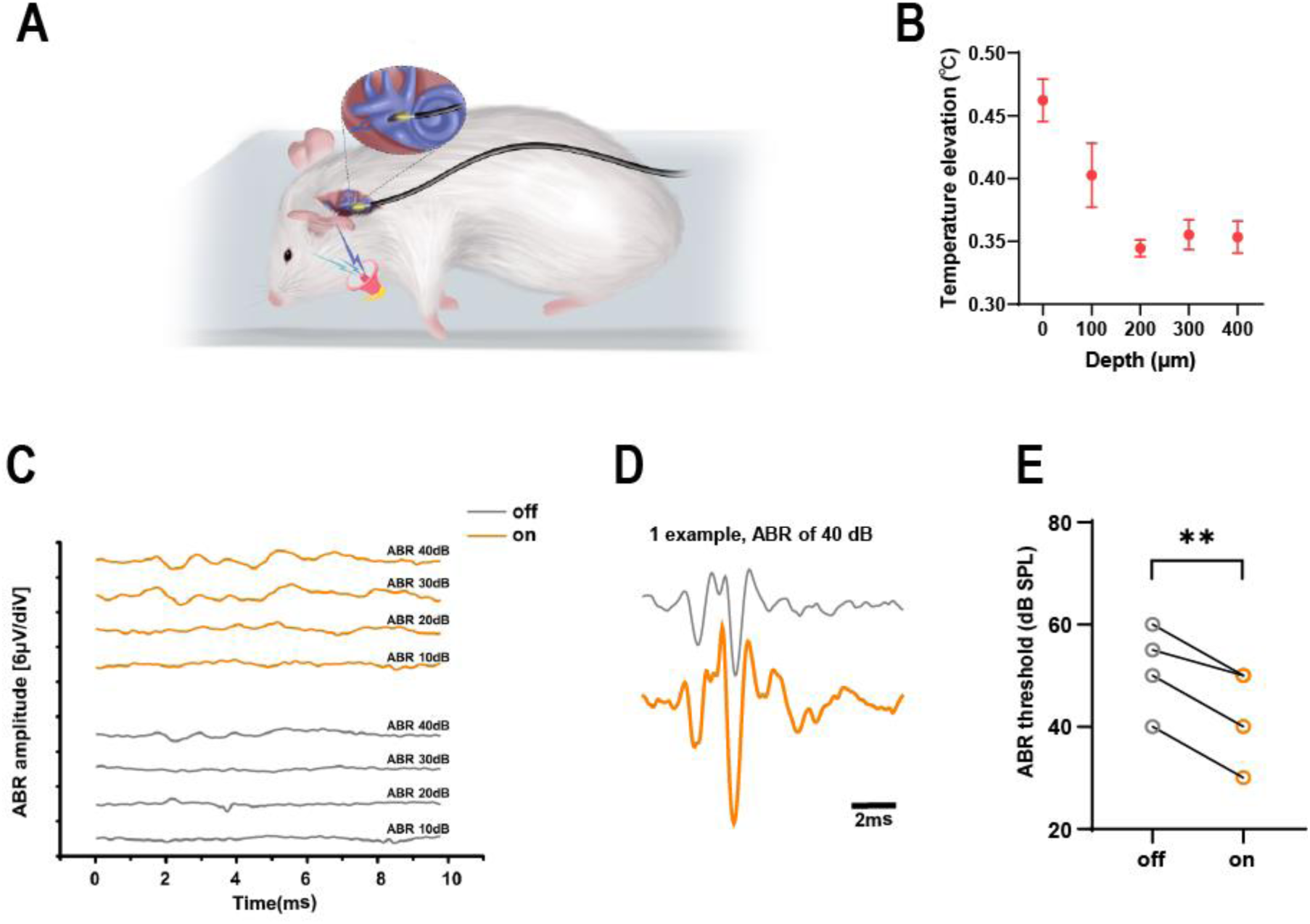
THM enhances guinea pig hearing perception. (**A**) Schematic drawing of the fenestration and ABR test in guinea pigs, together with the arrangements of THM. **(B)** Temperature elevation with respect to the distance from the fiber tip inside the cochlea. **(C)** A representative case showed the click ABR test in control (gray) and during THM (orange). A control guinea pig possessed a hearing threshold as low as 40 dB SPL. During THM, the guinea pig has 30 dB SPL hearing threshold at the same ear. **(D)** The start time of the ABR reaction did not change significantly, but the peak amplitude during THM was higher. **(E)** ABR threshold of guinea pig before and after irradiation.

A total of four animals were evaluated in the present study. Irradiation-induced temperature increase in the cochlea was below 0.5°C (Fig. 5B). The auditory brainstem response (ABR) signal was recorded in all four animals prior to cochlear fenestration, and click-sound stimuli were used at levels varying from 20 to 90 dB SPL. The ABRs all showed the classical Jewett waveform. The ‘control’ group did not receive THM, and the experimental group received THM in real-time during the listening test. Surprisingly, similar to our previous findings, guinea pigs in the THM treatment group exhibited an improved hearing threshold by an average of ∼10 dB (Fig. 5C). The peak value of ABR signals increased significantly, although the incubation period did not appear to change (Fig. 5D). When the click-sound stimulus was below their hearing threshold, the guinea pigs exhibited no ABR response, so OABR could be excluded. In summary, our results collectively indicate that THM affects sensorineural factors in guinea pigs, ultimately enhancing the sensitivity of their hearing perception (8.75 ± 1.25 dB, t3=7.000, P=0.006, n=4, Fig. 5E).

## Discussion

In this study, we found that THM significantly enhances the ability of MET channels in cochlear outer hair cells (OHCs) to respond to external mechanical stimuli. Furthermore, as irradiation power increases, the capability to respond also commensurately improves. In addition, THM increased the K^+^ currents without a significant effect on Na^+^ currents. Simulations of molecular dynamics showed that, through resonance, THM can adjust the efficiency of ion-selective filters of carbonyl-rich K^+^ channels. Cochlear hair cells convert sound stimuli into electrical signals by gating the mechanically sensitive ion channels in their cilia (hair) bundles (Furness et al., 2008), which is considered the first step in processing external sound information in the human ear (Gillespie et al., 2009).

After hair cells convert mechanical stimulation into electrical signals, K^+^ currents, as the predominant cochlear conductive current, contribute essential functions in the subsequent transmission and processing of electrical signals. Our results show that the application of THM significantly enhances both MET channels and K^+^ channels currents. Moreover, in the same recorded outer hair cells, different powers of THM all resulted in positive modulation of MET channel sensitivity (Fig. 1I), with the degree of enhanced sensitivity entirely dependent on the irradiation power, not on differences in external hair position of ciliated cells (Beurg et al., 2015). When the THM irradiation power is 75 mW, the MET current gain exceeds 50%, indicating that the OHCs are gain-modulated. OHCs can combine or integrate external mechanical stimuli through this mechanism to enhance auditory signals.

We further demonstrated that THM can increase the sensitivity of the hearing threshold in guinea pigs. In our experiment, THM irradiation led to an increase in the ABR amplitude of guinea pigs, but did not significantly change the incubation period. This finding led us to propose that the increase in hearing perception associated with THM treatment is therefore primarily sensorineural rather than conductive. Thus, THM at a specific wavelength can potentially enhance auditory function via neuromodulation to regulate signal processing at the molecular and cellular levels, and regulate sensorimotor processing at the behavioral level. We can thus easily envision the wide adoption of this method as a therapeutic strategy for sensorineural hearing loss.

Recent research into THM of ion channels has been generally based on carbon tube models, which have also been widely used to investigate the mechanisms of ion permeation (Wojciech et al., 2018; Johnson et al., 2011; Xiong et al., 2012). In order to clarify the molecular or atomic level biological effects of this treatment, we conducted MD simulations of ion permeation using a structural model of the ion channel protein containing the actual filter region determined by X-ray crystallography. Our results demonstrate that −C=O groups in cochlear K^+^ channels can resonantly absorb energy from THM, which then indirectly drives the collective vibration of −C=O groups, thus leading to an increase in K^+^ permeation through these channels. In contrast, no resonant wave absorption was detectable at 34.88 THz by −COO-groups in Na^+^ channels, nor was there any obvious change in Na^+^ permeation. Notably, the finger of the hydrogen bond network of water molecules is generally ranges from 0 ∼ 30 THz (Thrane et al., 1995), which effectively avoids 34.88 THz, indicating that the influence of THM is not due to thermal effects, but from the resonance absorption from THz waves to K^+^channels.

Previous studies have shown that near-infrared can produce OABR (Wang et al., 2016), and several mechanisms underlying the stimulation of nerve impulses by laser pulses have been proposed. For example, lasers can induce a thermal effect that activates thermionic ion channels in nerve cells of the spiral ganglia, leading to subsequent changes in the cell membrane and accompanying photomechanical effects that trigger the auditory nerve (Richard et al., 1971; Izzo et al., 2007). However, our experimental results show that instead of affecting spiral ganglion cells, THM can directly act on the cells that process the auditory signal one step further, the OHCs, consequently changing auditory sensitivity. Previous findings (Maréchal et al., 2011) indicate that the absorption rate of water is very low at the wavelength used in these experiments, therefore strongly suggesting that THM predominantly acts on functional molecules or proteins (such as MET channels and K^+^ channels) in a non-thermal manner.

In comparison with wearing hearing aids, stem cell differentiation and transplantation (Oshima et al., 2010; Li et al., 2003; Chen et al., 2012), optogenetics (Huet et al., 2021), and electronic cochlear implantation (Wilson et al., 1991; Kipping et al., 2020; Gang et al., 2008), THM requires no traumatic surgery, cumbersome equipment, or genetic manipulation, and is thus more suitable for use in human subjects. Our experiments also indicate that the effects of THM occur rapidly and reversibly within the timeframe of the experiment. Moreover, THM produces its regulatory effects on auditory signaling across relatively long distances, suggesting that modulation of the cochlear ion channels does not require contact between the light source and the tissue, preventing tissue damage and tissue-electrode compatibility problems. Given these potential advantages, THM warrants closer scrutiny as a promising method for therapeutic neuromodulation to regulate K^+^ channel-associated hearing disorders, brain functions and other diseases(Shi et al., 2020; Peng et al., 2020; Li et al., 2021).

## Materials and Methods

### Animals

C57BL/6J suckling rats (7 days old) and adult white guinea pigs (250-350g) were provided by the Experimental Animal Center of Tsinghua University. All experimental procedures were performed in accordance with the approval of the Animal Care and Use Committee of Tsinghua University and in accordance with institutional animal welfare guidelines.

### Mid-infrared light source

This research uses a quantum cascade Terahertz (THz) laser. The laser has a constant working wavelength of 8.6 μm. After the beam comes out, it is collimated by a beam collimator, and then the output is matched with the THz fiber (wavelength range: 3-17 μm, code: PIR600/700-100-FC/PC-FT-SP30, article number: AP12001, type: step index multimode) coupling, core diameter is 600 ± 15 μm, numerical aperture (NA) is 0.35 ± 0.05, and temperature range is -50 °C to +80 °C. The configuration is as follows: the pulse duration is 2 μs, the repetition frequency is 200 kHz, the duty cycle is 40%, and the current range is 0 mA to 1100 mA. The power is adjusted by changing the current.

### Cochlea anatomy

Mice were killed by decapitation. After removing the bony part and modiolus, dissected cochleae were transferred to a submersion-type chamber with anatomical fluid (142 NaCl, 6 KCl, 0.1 CaCl_2_, 1 MgCl_2_, 0.5 MgSO_4_, 3.4 L-glutamine, 8 D-glucose, and 10 H-HEPES, pH 7.4, osmolarity ∼290 mOsm/L, at room temperature, 21-23 °C). Cochleae were then mounted on an upright microscope, the attached muscle tissue and soft tissue were separated, volutes were peeled off, and the stria vascularis and tectorial membrane were torn in turn. Taking advantage of the adhesion of the vestibular membrane to the basement membrane, cochlear specimens were placed in a cell culture dish with extracellular fluid (144 NaCl, 0.7 NaH_2_PO_4_, 6 KCl, 1.3 CaCl_2_, 0.9 MgCl_2_, 5.6 D-glucose, and 10 H-HEPES, pH 7.4, osmolarity ∼320 mOsm/L, at room temperature, 21-23 °C), and with hair cells facing up.

### Whole-cell patch clamp

Before recordings, the petri dish containing the cochlear specimen was transferred into a recording room. Hair cells were visualized with an upright infrared differential interference contrast (IR-DIC) microscope (BX51WI, Olympus) equipped with a water-immersed objective (40x, NA 0.8). we performed whole cell recordings from the hair cells. For these recordings, patch pipettes were filled with a K^+^-based internal solution containing (in mM): 140 K-gluconate, 3 KCl, 2 MgCl_2_, 10 HEPES, 0.2 EGTA, and 2 Na_2_ATP (∼307 mOsm, pH 7.23 with KOH), and connected to a HEKA amplifier with a 5-kHz output filter. In order to check the properties of the MET channels, the hair bundles were stimulated on OHCs with a fluid jet applied through a glass electrode filled with bath solution (Johnson et al., 2011). Stimuli were applied using Patchmaster software (HEKA) and 20 psi air pressure as described previously (Xiong et al., 2012). Images were collected with a 1s sampling rate. Hair bundles were deflected with a glass pipette mounted on a P-885 piezoelectric stack actuator (Physik Instrument). The actuator was driven with voltage steps that were low-pass filtered at 10 kHz. Whole-cell recordings were carried out and currents were sampled at 100 kHz.

### Guinea pig fenestration surgery

In white adult guinea pigs (250-350 g), 2% sodium pentobarbital solution (0.15 mL/100 g) was injected intraperitoneally, and the operation was then performed after the animal was under anesthesia for 15 minutes. In order to ensure deep anesthesia during the experiment, the anesthesia state was judged by the plantar pinch reaction every 30 minutes and 0.1 mL of anesthetic was added as needed.

After the animal was depilated behind the ear, an arc-shaped incision was made, then a styloid-like structure was identified along the posterior wall of the external auditory canal to expose the auditory vesicle and the styloid-like structure. The muscle tissue and soft tissue attached to the styloid-like structure were removed, and a cranial drill was used to grind the posterior wall of the auditory vesicle. The opening was then expanded to expose the cochlea oval window, used for placement of optical fiber in subsequent THM experiments. During the experiment, the animals were placed on a 38 °C constant temperature platform to maintain a stable body temperature.

### Acoustic test and ABR recording

All sound stimuli in this study were written using BioSigRZ and generated using TDT System3 hardware (RP 2.1, PA 5, ED 1, HB 7). When calibrating the sound intensity, a microphone (Model 7016, ACO Pacific, Inc. USA) received the sound signal and converted it into an electrical signal, which was then collected by the TDT system and transmitted to the computer to determine the actual sound pressure level after calculation. The click-sound stimuli were used with varying levels from 20 to 90 dB SPL with a step size of 10 dB. In this study, unless otherwise specified, the closed sound field was used to play the sound. In this case, the front shielding sound and the detection sound were produced by the TDT classic field speaker EC 1, and transmitted sound to the animal’s ears through a thin tube of polymer material.

When measuring ABR, animals were first checked to ensure that they were still under anesthesia (additional injections of 0.1 mL 2% sodium pentobarbital solution in the intraperitoneal cavity were given as needed). After the animal was confirmed to be under anesthesia, the animal was placed on a soft cushion and the recording electrode, reference electrode and grounding electrode were placed. Three hypodermic needle electrodes were inserted into the middle point of both of the animal’s ears, behind the ears, and under the skin of the inner thighs, ensuring that the electrode resistance was less than 1KΩ. After the electrodes were properly placed, the recording started, and the ABR of the detection sound was recorded by TDT BioSigRZ and stored in the computer for online and offline analysis. The ABR sampling rate of the original signal was 25 kHz, and the filter setting during recording was 300 Hz for low pass and 3,000 Hz for high pass. During the experiment, the animal’s vital signs such as breathing, heartbeat and body temperature were monitored to ensure that the animal was in a normal state. Each individual stimulus was given 512 consecutive times to increase the signal-to-noise ratio on average. In the subsequent analysis, each individual wave of the ABR was recognized by the human eye and manually measured.

### Molecular dynamic simulation

Protein crystallization data demonstrates that K^+^ and Na^+^ channels are tetramers consisting of four single chains, including a narrow pore region (i.e., filter region), which have a decisive role in the permeation efficiency for K^+^ channel and Na^+^ channel. The critical factor to determine the permeability of K^+^ and Na^+^ concerns carbonyl (−C=O) groups and carboxyl (−COO^-^) groups arranged in the inner wall of the filter of K^+^ channel and Na^+^ channel, respectively (Chen et al., 2010; Noskov et al., 2004). The selective filter of K^+^ channel (PDB ID: 3LUT) is composed of 24 residues (sequence index 75∼80) (Chen et al., 2010), and the Na^+^ channel (PDB ID: 3RVY) has 40 residues (sequence index 1173∼1182) (Payandeh et al., 2011). Therefore, based on the function of the filter region, and in consideration of observing the comparative ion permeation events within the feasible time of molecular dynamics (MD), two simple models of K^+^ channel and Na^+^ channel were constructed (fig. S3). Then, the permeation of K^+^ and Na^+^ in a simulated physiological solution environment were compared. MD simulations were carried out at room temperature (300 K) without and with THM to compare the power of K^+^ and Na^+^ permeation for K^+^ channel and Na^+^ channel at nano-scale and femtosecond time resolution. The absorption spectra were calculated according to the Fourier transform of the velocity autocorrelation function of the total charge current of our simulation systems (Matthias et al., 2010). Based on the b3lyp/6-31g(d) method using Gaussian 09 software, the absorption mode corresponding to *γ* = 34.8 THz was identified by calculating the vibrations of −C=O groups for K^+^ channel. The b3lyp/6-31g(d) method was then used to explore the vibration spectra of the filter region containing 3 K^+^ for K^+^ channel. The filter region was simplified to form the one-dimensional (1D) chain of O^2-^-K^+^-O^2-^-K^+^ containing O^2-^ and K^+^ (inset of Fig. 4C), and then the collective vibrations of K^+^ connected with the −C=O groups were calculated based on the linear lattice model. For details, refer to section 1.3 of Supplementary Information (SI).

### Light spot test

The laser beam analyzer (Swiss RIGIM series, M2: 640 × 480) and e-bus software were used to measure the characteristics of the beam output after the beam passed through the collimator and optical fiber. During the measurement, the output of the laser was set to the same output mode as in the experiment, and the distance between the output end surface and the test surface of the beam analyzer was 1mm. The obtained light spot characteristic diagrams are expressed in cross-sectional mode (fig. S1A) and three-dimensional mode (fig. S1B). The optical spot output by the optical fiber is approximated as a Gaussian spot. According to the rule, the distance from the center point when the laser light intensity was reduced to 1/e^2^ of the peak was defined as the spot radius, the measured spot radius was 2 mm.

### Temperature measurement

We followed the tissue temperature measurement protocols as in the previous literature25. The sensor (MT 29/5, Physitemp) was calibrated once before the experiments and the distance between the sensor and the top of the fiber is controlled by a stereotactic locator under the microscope.

### Data analysis

Data analysis was performed using MATLAB (MathWorks, Bethesda, MD) and Igor pro 8 (WaveMetrics, Lake Oswego, OR). The sample size was not predetermined, but was based on previously reported figures. All measurements were taken from different samples. To compare conditions, data from at least six cells were obtained from different animals. No samples were excluded from the analysis. The data are expressed as mean ± s.e.m. The error bars in the figures also represent s.e.m. Unless otherwise specified, if the data are paired, a two-tailed paired Student’s *t*-test was used. For two independent observations, an independent Wilcoxon rank-sum test was used to compare abnormal data. For more than two independent observations, repeated analysis of variance and Bonferroni’s multiple comparisons were used. If P<0.05, the difference was considered significant.

## Acknowledgments

This work was supported by the National Defense Science and Technology Innovation Special Zone, and the National Supercomputer Center in Tianjin. C.C. acknowledges the support from the XPLORER Prize.

## Supplementary figures and legends

### 1.1 The light spot

**Figure. S1:**
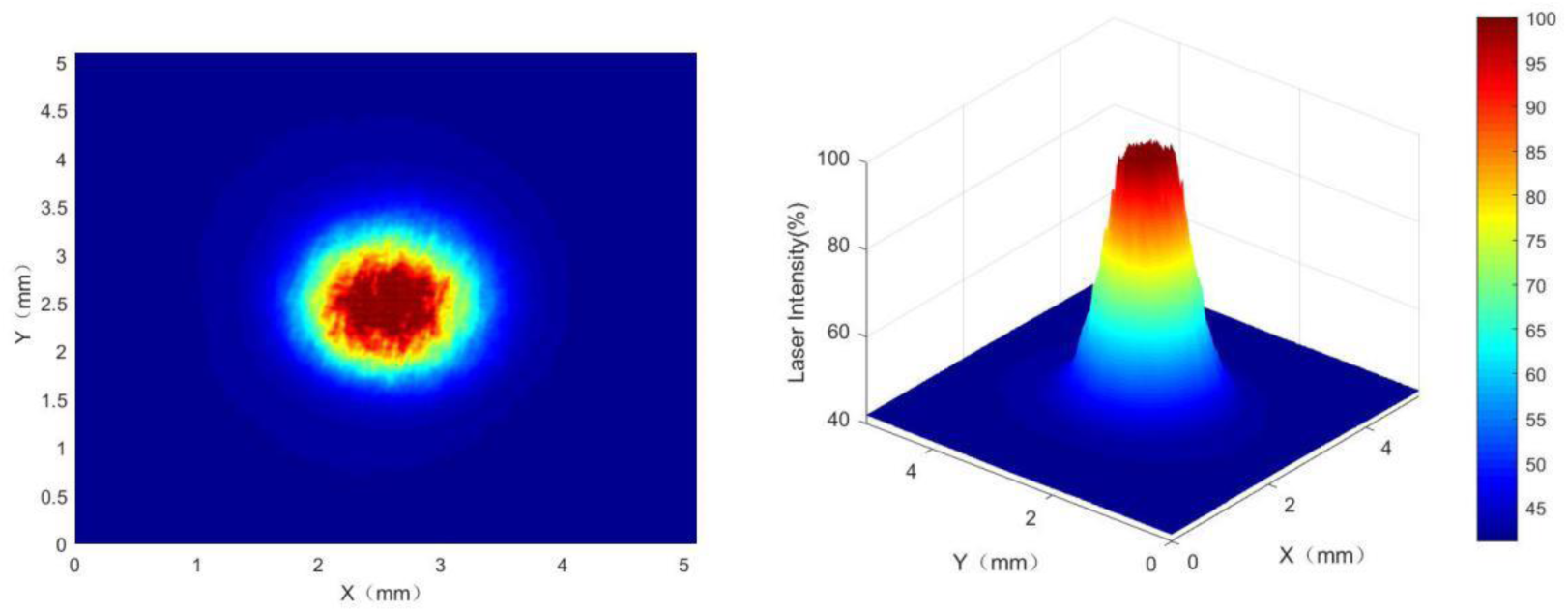
At a working distance of 1 mm, the output light spot characteristics after the beam is collimated. Left: Cross-sectional view; Right: three-dimensional view.

### 1.2 Temperature variation

**Figure. S2:**
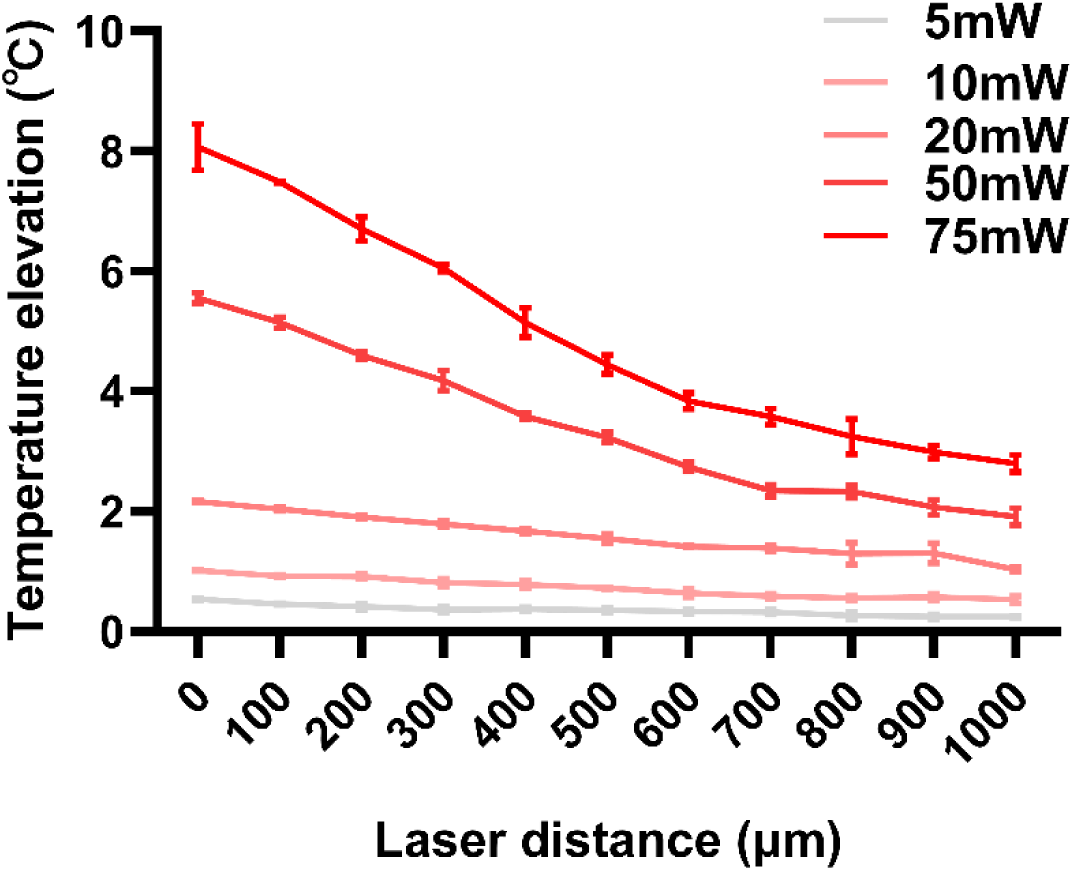
Temperature variation with distance at different power (n=6).

### 1.3 The structure model

In the calculation model, the K^+^ channel (PDB ID: 3LUT)^1^ was constructed containing the filter region, and carbon tubes were placed at z = 2.412 nm, z = 3.712 nm in the middle to separate water and ions on each side. There were 12 K^+^, 12 Na^+^, and 20 Cl^-^ on each side, respectively, of the carbon tube to generate the potential barrier on the K^+^ channel. Thus, the wall setup (materials of C–C type) was ∼ 0.3 nm thickness, located at z=0, z=z_box. The total charge was electrically neutral (Fig. S.3a). Similarly, for Na^+^ channel (PDB ID: 3RVY)^2^, the filter region was constructed, and carbon tubes were placed at z = 3.494 nm and z = 4.606 nm in the middle positions. There were 14 Na^+^, 14 K^+^, and 24 Cl^-^ on both sides of the carbon tube, and the wall setup (materials of C–C type) also had ∼ 0.3 nm thickness, located at z = 0, z = z_box. The total charge was also electrically neutral (Fig. S.3b).

Considering the effects of THM on the ion permeation process, THM with frequency *γ* = 34.88 THz was added into the whole simulation system according to the formula, ***E***(t) = *A*·**u**·cos(ωt + φ), where *A* represents the maximum amplitude of the electric field, which determines the strength of electric field component of electromagnetic wave; and **u** and phi represent the polarization direction and phase, which are set to (0, 0, 1) and 0, respectively^3,4^. The electromagnetic wave frequency γ is related to the angular frequency ω by the equation γ = ω/2π.

Based on molecular dynamics package GROMACS^5^, the ion permeation with THM was simulated at 300 K. The simulation process can be divided into two stages. First, the initial simulation system was minimized to eliminate unreasonable conformation. Then, under the NVT ensemble, both the carbon tube and filter region (except –C=O groups for K^+^ channel, –COO^-^ groups for Na^+^ channel) in the system were fixed, and the –C=O groups for K^+^ channel, –COO^-^ groups for Na^+^ channel of the filter region were released freely. Based on these above procedures, THM with characteristic frequency *γ* = 34.88 THz was applied, the total MD simulation time was set as ∼ 1 μs, and the trajectory was saved every 0.1 ps for subsequent data analysis. In the simulation, the force field of Charmm 27^6^ and periodic boundary conditions were used. The connection element Ewald sum algorithm^7^ was used to deal with the electrostatic interaction. In the simulation process, all bond lengths were limited by the Lincs algorithm^8^. Then, the Velocity-Verlet algorithm^9^ was performed to solve the atomic motion equation, and the time step was set as 2 fs. In particular, for the K^+^ channel, the truncation of the Lennard-Jones interaction and the real space part of the Ewald sum is 1.90 nm, the convergence factor of the Ewald sum is 1.65 nm, and the radius of the K-space section is 10.4 nm. Similarly, for the Na^+^ channel, the truncation of Lennard-Jones interaction and the real space part of Ewald sum is 1.623 nm, Ewald and convergence factor are 1.65 nm, and the radius of the K-space section is 10.4 nm.

The absorption spectra were calculated based on the Fourier transform of the autocorrelation function of the total charge current *J*(t) = Σ_i_q_i_v_i_(t)^10^, where q_i_ represents the charge of the i-th atom, and v_i_(t) stands for the velocity of the i-th atom at time t.

**Figure. S3:**
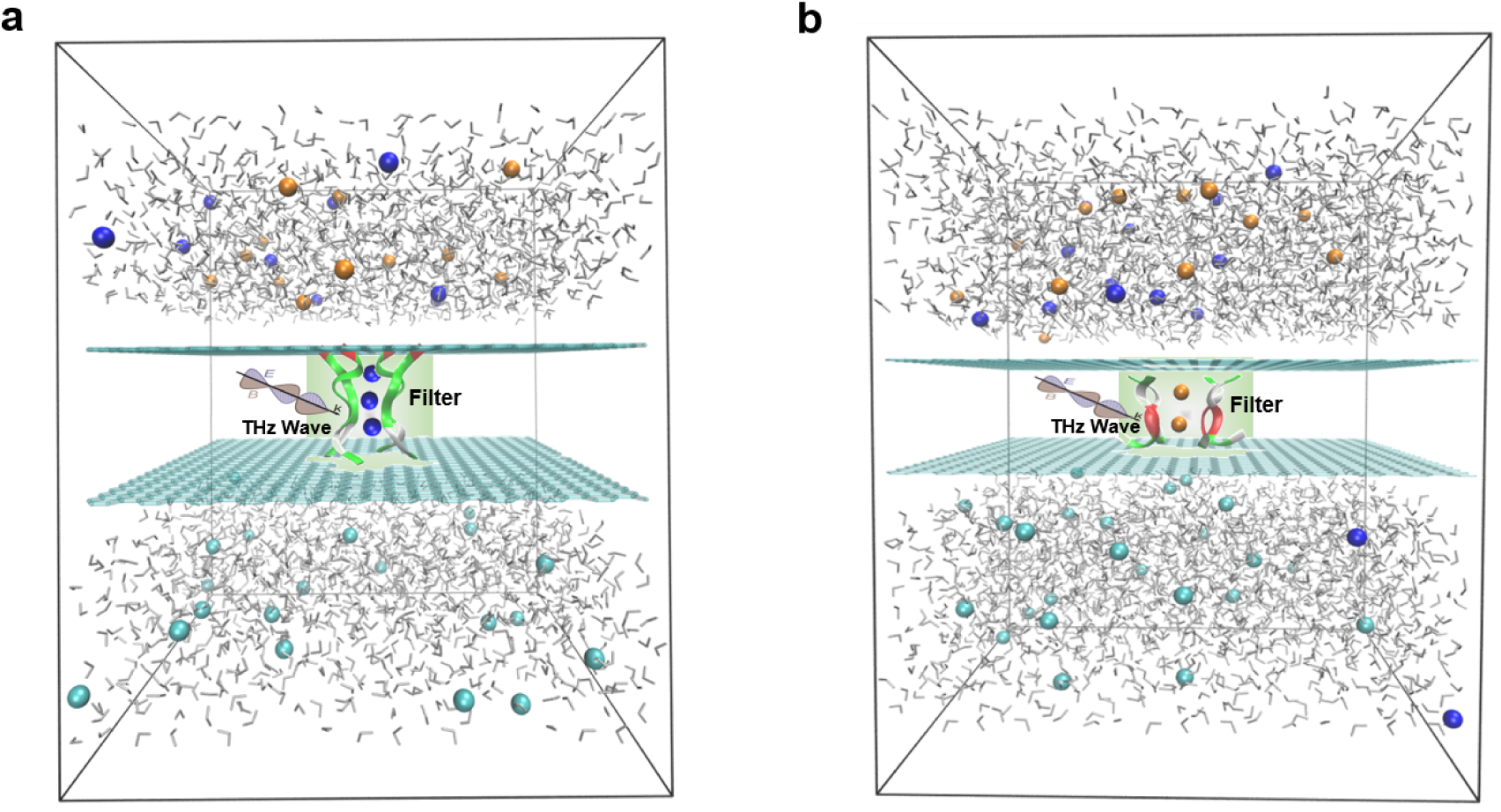
The atomic models of the K^+^ channel and Na^+^ channel. The K^+^ channel (PDB ID: 3LUT)^1^ model contains 10,294 atoms, including 2,725 TIP3P water molecules, 12 K^+^, 12 Na^+^, and 20 Cl^-^. The protein has four negative charges, the addition of four counterbalance ions ensures the whole system is electrically neutral, and thus the concentration of a salt solution is 0.15 M. The size of the PBC box is 5.04 nm × 5.16 nm × 6.25 nm. **a** The Na^+^ channel (PDB ID: 3RVY)^2^ model contains 14,922 atoms, including 4,067 water molecules, 14 Na^+^, 14 K^+^, and 24 Cl^-^. Similarly, the addition of four counterbalance ions ensures the whole system is electrically neutral, so that the concentration of a salt solution is 0.15 M. The size of the PBC box is 5.40 nm × 5.40 nm × 7.30 nm. The blue atom represents K^+^, the yellow atom represents Na^+^, the green atom represents Cl^-^, and the TIP3P water molecule is displayed in the format of a ball stick **b**.

### 1.4 Eigen-mode of –C=O vibrations for K^+^ channel

**Figure. S4:**
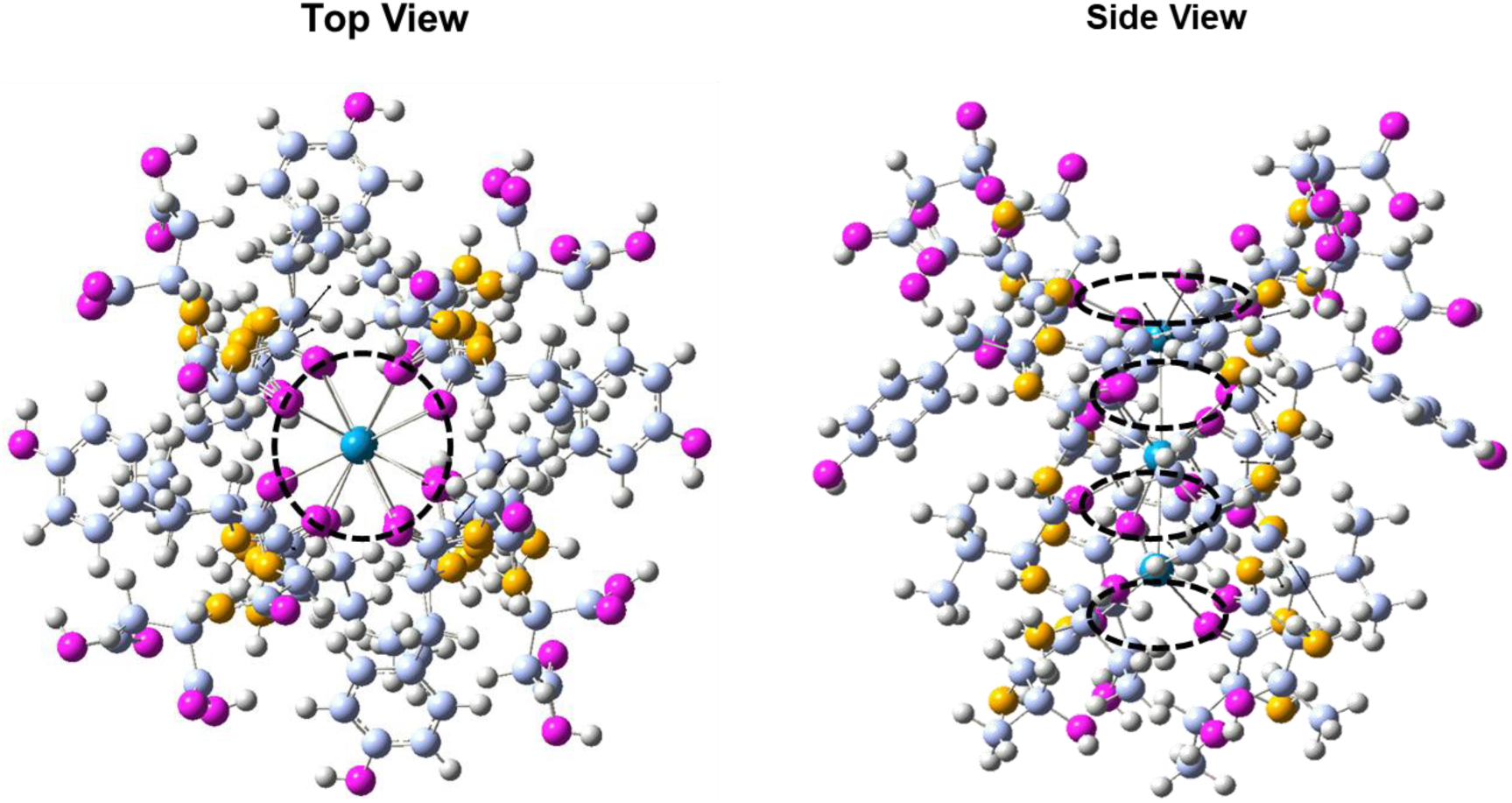
The collective oscillation mode of carbonyl (–C=O) groups corresponding to the frequency *γ* = 34.88 THz for the K^+^ channel based on Gaussian 09 software. The section in the circle from the top view, shows that the distances among four groups of – C=O connections from the center of the system are equal or close to equal. From the side view, the four groups of –C=O vibrations are basically in the same plane.

### 1.5 The linear lattice model

The equation of motion of K^+^ confined in 1D K^+^ channel was written as (*11*),

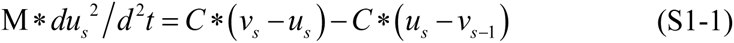

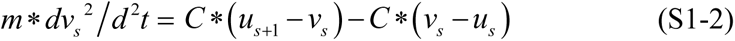

where *M* and *m* indicate the masses of O^2-^ and K^+^, respectively; *s* denotes the label of a unit cell; *u* and *v* are the displacements of O^2-^ and K^+^ near each lattice site, respectively; and *C* is the coupling constant between the nearest neighboring atoms. The function form of 1D lattice wave was given as^11,^ 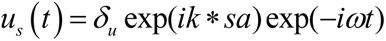 (S2-1), 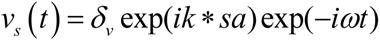 (S2-2) where *δ_u_* and *δ_v_* represent maximum amplitudes of the displacement for O^2-^ and K^+^ ions. By substituting formula (2) into formula (1), we then derived the results of the formula, which are shown in equation 1.

